# Differential repression of *Otx2* underlies the capacity of NANOG and ESRRB to induce germline entry

**DOI:** 10.1101/2021.06.14.448276

**Authors:** Matúš Vojtek, Jingchao Zhang, Juanjuan Sun, Man Zhang, Ian Chambers

## Abstract

Primordial germ cells (PGCs) are induced in the embryo by signals, including BMP emanating from extra-embryonic ectoderm, that act on cells in the post-implantation epiblast. PGC development can be recapitulated *in vitro* through the exposure of epiblast-like cells (EpiLCs) to appropriate cytokines, resulting in differentiation into PGC-like cells (PGCLCs). Interestingly, the requirement for cytokines to induce PGCLCs can be bypassed by enforced expression of the transcription factor (TF) NANOG. However, the underlying mechanisms are not fully elucidated. Here, we show that *Otx2* downregulation is essential to enable NANOG to induce PGCLC formation. Moreover, while previous work has shown that the direct NANOG target gene Esrrb can substitute for several functions of NANOG, enforced expression of ESRRB cannot promote PGCLC specification from EpiLCs. This appears to be due to differential downregulation of *Otx2* by NANOG and ESRRB, since induction of ESRRB in *Otx2^+/-^* EpiLCs activates expression of the core PGC TFs, *Blimp1, Prdm14* and *Ap2*γ and emergence of PGCLCs. This study illuminates the interplay of TFs occurring at the earliest stages of PGC specification from a state of competence.

## Introduction

Germline development and sexual reproduction depend on the establishment of primordial germ cells (PGCs). In mice, PGCs are induced by bone morphogenic factor 4 (BMP4) acting on proximal posterior epiblast cells at embryonic day 6 (Lawson et al., 1999). Prospective PGCs downregulate the epiblast transcription factor (TF) *Otx2* and subsequently activate the key PGC TFs Blimp1 (Prdm1), Prdm14 and Ap2γ (Kurimoto et al., 2008; Ohinata et al., 2005; Vincent et al., 2005; Weber et al., 2010; Yamaji et al., 2008; Zhang and Chambers, 2019). These events can be recapitulated *in vitro* by differentiating naïve embryonic stem cells (ESCs) into epiblast-like cells (EpiLCs), which are transiently competent to specify PGC-like cells (PGCLCs) in response to BMP4 and associated cytokines (Hayashi et al., 2011; Hayashi and Saitou, 2013). The requirement for BMP4 and associated cytokines can however be bypassed by induction of NANOG in EpiLCs (Murakami et al., 2016). Combined ectopic expression of NANOG and cytokine treatment further expands the proportion of PGCLCs in the differentiated population (Murakami et al., 2016). Even though Nanog is not essential for emergence of PGCs, nor for germline transmission (Carter et al., 2014; M. Zhang et al., 2018), *Nanog* deletion results in a large decrease in PGC numbers both *in vitro* and *in vivo* (Chambers et al., 2007; Murakami et al., 2016; Yamaguchi et al., 2009; M. Zhang et al., 2018).

The target genes through which NANOG acts in ESCs have been identified and include *Esrrb* and *Otx2*, which are regulated positively and negatively by NANOG, respectively (Festuccia et al., 2012). Both of these genes also regulate the PGC compartment (Mitsunaga et al., 2004; J. Zhang et al., 2018). *Esrrb* can rescue several functions of NANOG in *Nanog*-null pluripotent cells (Festuccia et al., 2012). Interestingly, analysis of the germline indicates that loss-of-function for *Esrrb* results in a similar quantitative reduction in PGC numbers at midgestation as observed when *Nanog* is deleted specifically from the germline (Mitsunaga et al., 2004; M. Zhang et al., 2018). Moreover, deletion of *Nanog* impairs induction of PGCLC differentiation in response to PGC-promoting cytokines (Murakami et al., 2016). This absence of PGC differentiation in response to cytokines can be compensated for by enforced expression of ESRRB (M. Zhang et al., 2018). Consistent with a conserved epistatic relationship between Nanog and Esrrb both in the preimplantation epiblast and in the germline, a knock-in of *Essrb* to the *Nanog* locus overcomes the reduction in PGC numbers resulting from germline specific *Nanog* deletion (M. Zhang et al., 2018). In ESCs, Otx2 and Nanog antagonise each other by mutual repression (Acampora et al., 2017; Festuccia et al., 2012). We have shown that entry to the germline is blocked when OTX2 expression is maintained during the first two days of EpiLC - PGCLC differentiation (J. Zhang et al., 2018; Zhang and Chambers, 2019). In contrast, *Otx2* deletion dramatically increases PGCLC numbers *in vitro* and raises PGC numbers *in vivo* (J. Zhang et al., 2018). Recently, OTX2 has been shown to act through cis-acting binding sites that repress transcription of *Nanog* and *Oct4* (Di Giovannantonio et al., 2021). However, despite an increasing knowledge of the relationships between Esrrb, Nanog and Otx2, the interplay between these factors in PGC induction is not completely understood. Here we examine the capacity of ESRRB and NANOG to induce PGCLC differentiation using inducible transgene systems in the absence of cytokines. Our results uncover a differential capacity of NANOG and ESRRB to repress *Otx2* and an OTX2 dose-dependent barrier to germline induction by ESRRB and NANOG in the absence of cytokines.

## Results

### Esrrb cannot activate PGCLC programme

To examine whether ESRRB can induce cytokine free PGCLC differentiation similarly to NANOG, we overexpressed NANOG or ESRRB during cytokine free PGCLC differentiation. Cell lines carrying tetracycline inducible Nanog (TgiN) or Esrrb (TgiE) transgenes were generated by integrating piggybac transposons into E14Tg2a ESCs expressing the modified reverse tetracycline transactivator (rtTA2) (Urlinger et al., 2000) from *Rosa26* (Figure 1A). The resulting TgiN and TgiE ESCs, cultured in 2i/LIF medium, were then differentiated to PGC-competent epiblast-like cells (EpiLCs) by two-day culture in the presence of Activin A and bFGF (Figure 1B). In the original PGCLC differentiation protocol, EpiLCs are aggregated and stimulated using a cytokine cocktail that is required for PGCLC specification (Hayashi et al., 2011; Hayashi and Saitou, 2013). However, NANOG has been shown to direct PGCLC differentiation in the absence of cytokines (Murakami et al., 2016). We therefore omitted cytokines and tested the ability of NANOG or ESRRB overexpression to induce PGCLC development (Figure 1B). TgiN and TgiE cells formed similar colonies both as naïve ESCs and EpiLCs (Figure S1A). Addition of doxycycline during PGCLC specification from EpiLCs activated robust induction of Nanog or Esrrb transgenes in TgiN or TgiE cells, respectively (Figure S1B). In the absence of doxycycline, TgiN and TgiE cells did not express the transgenes (Figure S1B). Induced expression of either Nanog or Esrrb resulted in similar levels of surface expression of SSEA1 and CD61 which were considered to jointly mark PGCLCs (Hayashi et al., 2011) (Figure 1C, D). However, in contrast to NANOG, induction of ESRRB failed to increase the expression of *Blimp1 or Prdm14* mRNAs and showed only a modest increase in *Ap2γ* mRNA level (Figure 1E). We therefore assessed changes in PGC TF expression earlier, during the first 48 hours of differentiation (Figure 1F, S1B). Induction of Nanog activated expression of both *Esrrb* and the PGC transcription factors *Blimp1, Prdm14* and *Ap2γ*. Interestingly, while induction of ESSRB increased *Blimp1, Prdm14* and *Ap2γ* mRNAs within the first 6 hours of differentiation, ESRRB failed to sustain *Blimp1* and *Prdm14* expression at later time points (Figure 1F). Therefore, ESRRB unlike NANOG does not produce a sustained activation of the PGC programme during cytokine-free PGCLC differentiation.

**Figure 1:**
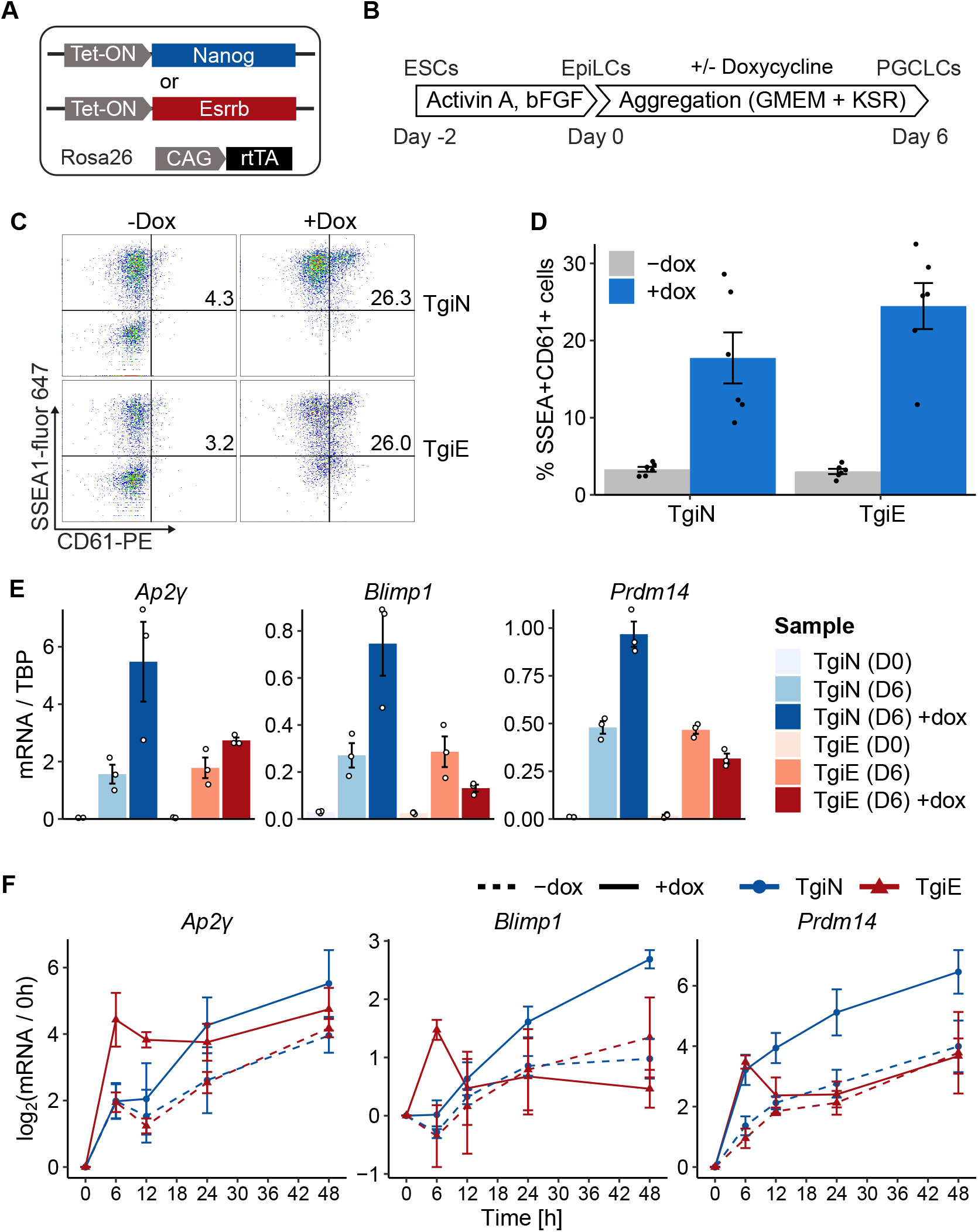
Esrrb cannot activate the PGC programme. **A)** Cartoon of TetON-Nanog (TgiN) and TetON-Esrrb (TgiE) cell lines. TetON-Nanog and TetON-Esrrb expression cassettes were randomly integrated into E14Tg2a cells expressing rtTA from *Rosa26*. **B)** The cytokine-free PGCLC differentiation protocol. ESCs cultured in 2i/LIF were differentiated to EpiLCs by culture in Activin A and bFGF for 2 days. EpiLCs were then aggregated in GMEM+KSR medium for 6 days in the presence or absence of doxycycline and PGCLC status was analysed. **C)** Representative example of flow cytometry analysis of PGCLC aggregates from TgiN and TgiE cells at day 6 with or without doxycycline, showing the percentage of SSEA1^+^CD61^+^ cells. **D)** Quantification of **C**; error bars are mean ± SEM, n = 6. **E)** RT-qPCR quantification of indicated transcripts in TgiN and TgiE cells at day 6 of cytokine-free PGCLC differentiation in the presence (+dox) or absence (-dox) of doxycycline. Bars represent mean mRNA levels normalised to TBP mRNA; points are individual data measurements; error bars are mean ± SEM, n = 3. **F)** Time course analysis of the indicated mRNAs during the first 48 hours of cytokine-independent PGCLC differentiation of TgiN and TgiE EpiLCs with (+dox) or without (-dox) doxycycline. Points, triangles and lines represent mean log2 fold change of normalised mRNA levels over zero-hour time point (mean ± SD; n = 3).

### Nanog induces PGCLCs by repressing *Otx2*

Our previous results show that the requirement of cytokines for PGCLC formation is also eliminated in *Otx2^-/-^* cells (J. Zhang et al., 2018). OTX2 and NANOG have antagonistic functions in ESCs (Acampora et al., 2017). Moreover, NANOG can directly downregulate *Otx2* in ESCs (Figure 2A; Festuccia et al., 2012; Heurtier et al., 2019). Microarray data suggests that this capacity to repress *Otx2* may not be shared by ESRRB (Figure 2A; Festuccia et al., 2012). This raises the hypothesis that ESRRB cannot effectively induce PGCLC specification due to an impaired capacity to repress *Otx2*. To address this hypothesis, we first measured *Otx2* mRNA expression within the first 48 hours of EpiLC aggregation of TgiN and TgiE cells in cytokine-free medium (Figure 1B). NANOG induction drove a rapid downregulation of *Otx2* between 6 and 12 hours compared to uninduced cells (Figure 2B) agreeing with the hypothesis that NANOG represses *Otx2* during PGC specification. In contrast, ESRRB induction did not affect *Otx2* mRNA levels during the first 24 hours and even showed increased *Otx2* mRNA levels at 48 hours (Figure 2B).

**Figure 2:**
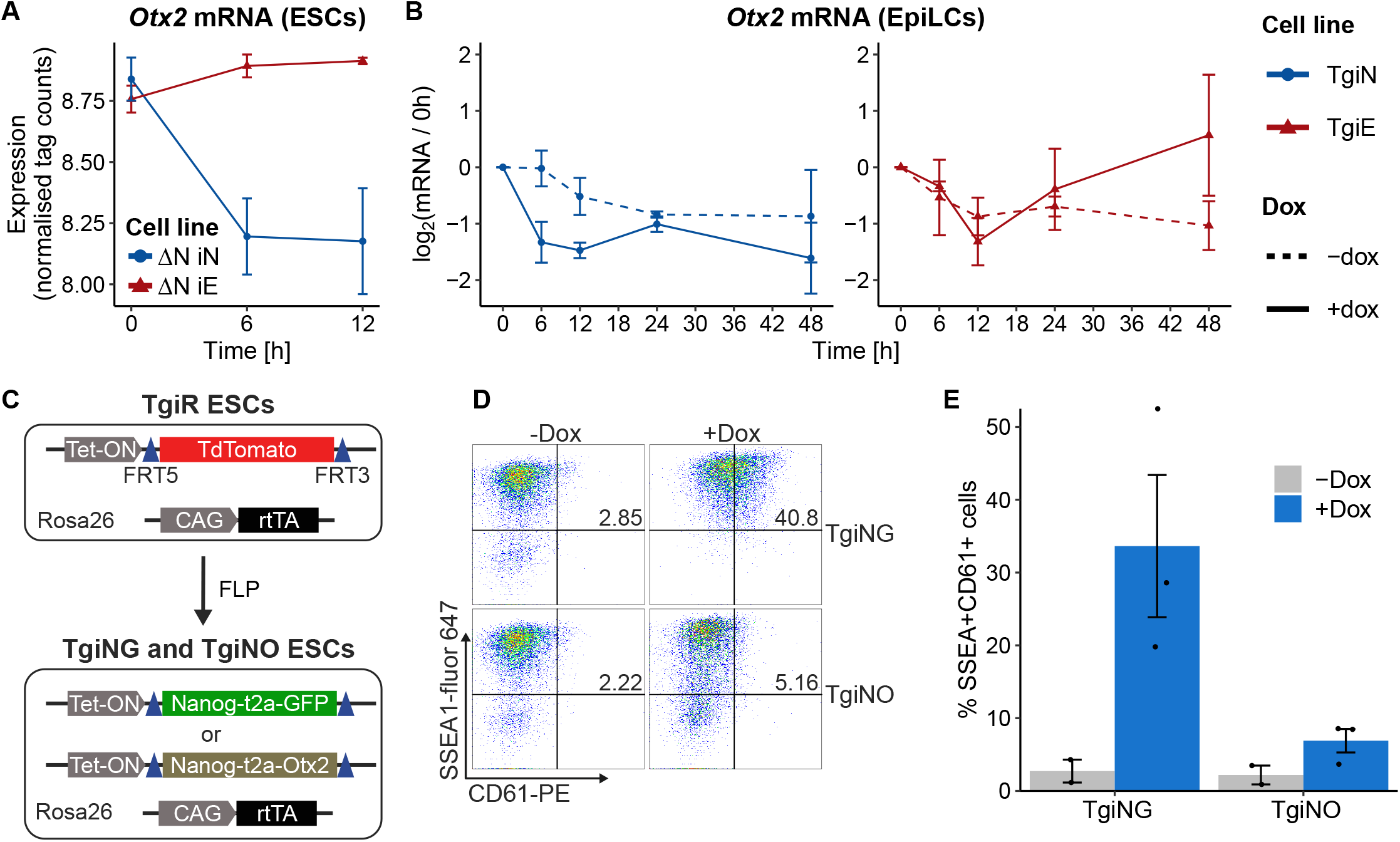
Otx2 downregulation is essential for PGCLC induction by Nanog. **A)** Relative changes of *Otx2* mRNA levels in *Nanog^-/-^* TetON-Nanog (ΔN-iN) or *Nanog^-/-^* TetON-Esrrb (ΔN-iE) ESCs at indicated timepoints after doxycycline treatment (mean ± SD, n = 3). Data adapted from (Festuccia et al., 2012). **B)** Relative changes of *Otx2* mRNA levels after aggregation of doxycycline inducible TgiN and TgiE EpiLCs cultured in the absence (-dox) or presence (+dox) of doxycycline. Lines, points and triangles represent mean log2 foldchange (FC) differences between data points and the zero-time timepoint (mean ± SD; n=3). **C)** *Rosa26:* rtTA; E14Tg2a-TetON-TdTomato (TgiR) ESCs were modified as shown to derive inducible Nanog-t2a-GFP (TgiNG) or Nanog-t2a-Otx2 (TgiNO) ESCs from. **D)** Representative example of flow cytometry analysis of PGCLC aggregates from TgiNG and TgiNO cells at day 6 with or without doxycycline, showing the percentage of SSEA1^+^CD61^+^ cells. **E)** Quantification of **D** showing percentage of SSEA-1^+^/CD61^+^ cells (mean ± SEM; n = 2 for -Dox, 3 for +Dox).

We next tested whether OTX2 clearance is necessary for NANOG to induce PGCLCs. To do this, we first generated E14Tg2a ESC lines which can induce either GFP (TgiNG) or OTX2 (TgiNO) from the same transgene that induces NANOG (Figure 2C). The TgiNG and TgiNO cells were generated by replacing the TdTomato coding sequence in E14Tg2a TetON-TdTomato (TgiR) cells with cassettes encoding Nanog-t2a-GFP or Nanog-t2a-Otx2 (Figure 2C). While doxycycline has no effect on *Nanog* or *Otx2* expression in the parental TgiR cell line (Figure S2), doxycycline treatment increased *Nanog* expression ~8 and 5-fold in TgiNG and TgiNO cells, respectively (Figure S2). In addition, *Otx2* mRNA was induced by doxycycline only in TgiNO cells (Figure S2). To assess the effect of OTX2 on the ability of NANOG to induce PGCLC differentiation, TgiNG and TgiNO cells were subjected to PGCLC differentiation in the absence of cytokines. Simultaneous induction of GFP and NANOG upregulated expression of PGCLC surface markers CD61 and SSEA1 (Figure 2D, E). In contrast, simultaneous induction of OTX2 and NANOG from the same doxycycline-responsive transgene markedly reduced the population of SSEA1^+^/CD61^+^ cells (Figure 2D, E). This indicates that the capacity of NANOG to function in PGCLC induction requires repression of *Otx2*.

### Otx2 heterozygosity enables Esrrb to induce cytokine-free PGC differentiation

To test whether *Otx2* is a limiting factor that prevents ESRRB from activating PGCLC specification, we integrated doxycycline inducible Nanog (iN) or Esrrb (iE) transgenes into heterozygous *Otx2*^lacZ/fl^ ESCs (Acampora et al., 2013; J. Zhang et al., 2018) (Figure 3A). This cell line also contains a GFP transgene which reports the activity of the Oct4 distal enhancer (ΔPE::GFP) (Figure 3A) which becomes activated in PGCLCs (Magnusdottir et al., 2013). We refer to these cells as *Otx2^+/-^* iE and *Otx2^+/-^* iN cells. We isolated two *Otx2^+/-^* iE clones (1 and 10) and one *Otx2^+/-^* iN clone that each express approximately 50% the level of *Otx2* mRNA compared to *Otx2*^+/+^ cells in the EpiLC state (Figure 3B).

**Figure 3:**
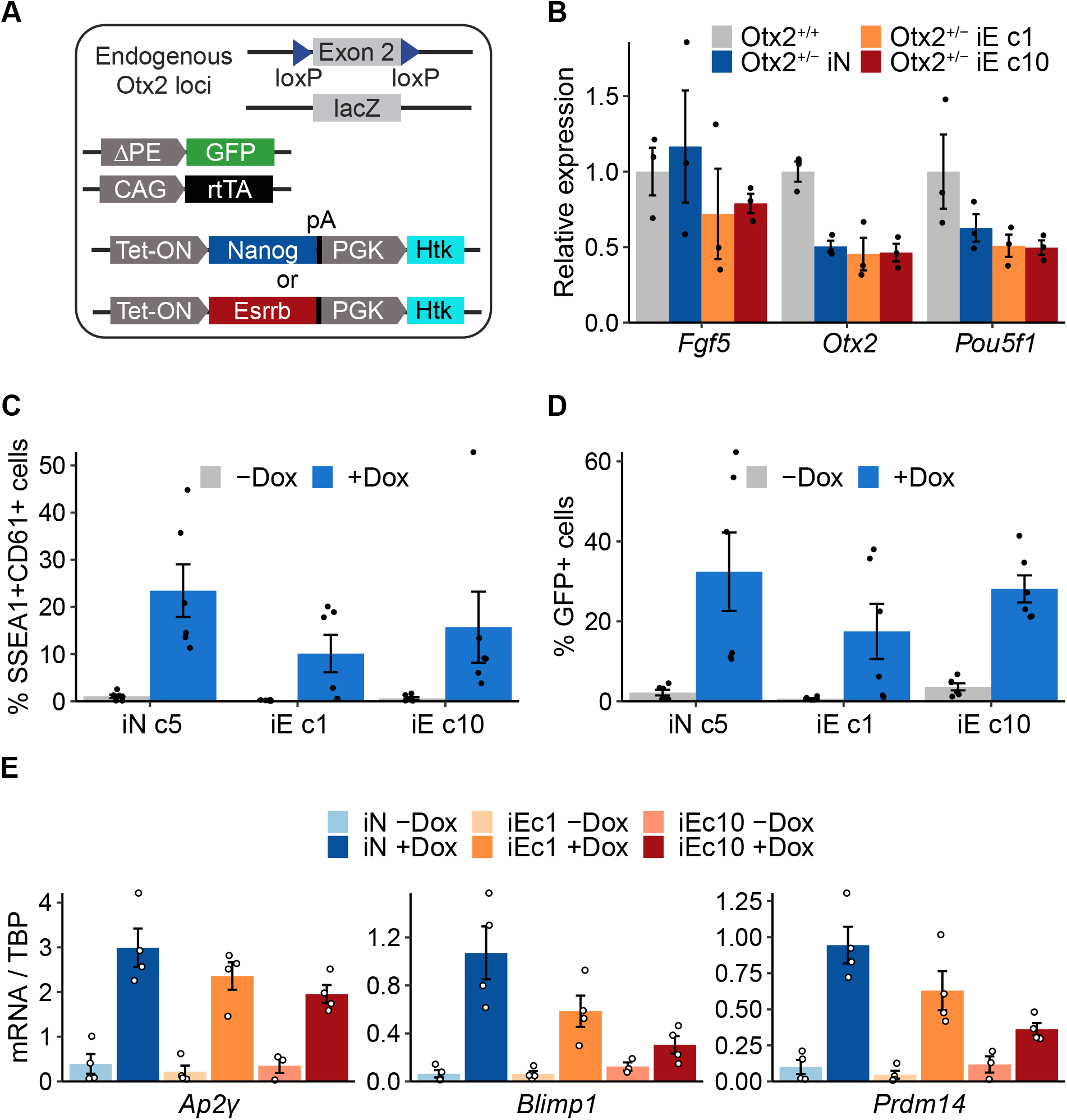
A reduced *Otx2* gene dose facilitates cytokine-free germline entry by Esrrb induction. **A)** Diagram of *Otx2^+/-^: Oct4* ΔPE-GFP ESCs carrying doxycycline inducible Nanog (iN) or Esrrb (iE) transgenes. **B)** Expression levels of the indicated mRNAs in *Otx2^+/-^* iN and iE (c1 and c10) EpiLCs in the absence of doxycycline. Bars are the mean mRNA levels, normalised to TBP mRNA; points are individual data measurements; error bars are mean ± SEM, n = 3. **C)** Size of PGCLC population in the presence (+Dox) or absence of (-Dox) doxycycline at day 6 of cytokine-free PGCLC differentiation of the indicated Otx2^+/-^ EpiLCs represented by SSEA1 and CD61 co-expression, mean ± SEM; n = 6). Individual measurements are shown. **D)** Proportion of cells expressing the *Oct4* ΔPE-GFP reporter in the indicated *Otx2^+/-^* aggregates at day 6 of the cytokine-free PGCLC differentiation. Bars represent mean ± SEM of percentage of GFP+ cells (n = 6). **E)** RT-qPCR analysis of the indicated mRNAs in day 6 aggregates from **C**. Bars are means ± SEM (n= 4).

Doxycycline treatment of *Otx2^+/-^* iN and *Otx2^+/-^* iE cell lines increased expression of the transgenes by 20 to 40-fold compared to wild-type E14Tg2a cells at day 2 of cytokine-free PGCLC differentiation (Figure S3A). When *Otx2^+/-^* cell lines were subjected to cytokine-free PGCLC induction in the absence of doxycycline, surface expression of CD61/SSEA1 was not induced in iE or iN cell lines (Figure S3B). However, upon induction of either NANOG or ESRRB by doxycycline, surface expression of CD61/SSEA1 was induced (Figure 3C, S3B). In addition, the proportion of cells expressing the Oct4 ΔPE::GFP transgene was similarly induced in each of these lines by doxycycline treatment (Figure 3D). Furthermore, induction of either ESRRB or NANOG in *Otx2^+/-^* cells increased expression of *Blimp1, Prdm14* and *Ap2γ* mRNAs at day 6 (Figure 3E). This contrasts with induction of ESRRB in *Otx2^+/+^* cells which did not increase *Blimp1* and *Prdm14* mRNA levels (Figure 1E). Interestingly, at the earlier time point of day 2 of differentiation, ESRRB induced lower levels of *Blimp1, Prdm14* and *Ap2γ* mRNA expression in *Otx2^+/-^* cells than were achieved by NANOG induction (Figure S3A). This may indicate that ESRRB-induced germline entry in *Otx2^+/-^* cells is delayed compared to that induced by NANOG.

Based on these results we propose a model in which NANOG induces germline fate predominantly by downregulation of *Otx2* (Figure 4). Once OTX2 levels are substantially diminished, NANOG and its direct downstream target ESRRB can activate PGC determinants *Blimp1, Prdm14* and *Ap2γ* and initiate germline specification (Figure 4).

**Figure 4:**
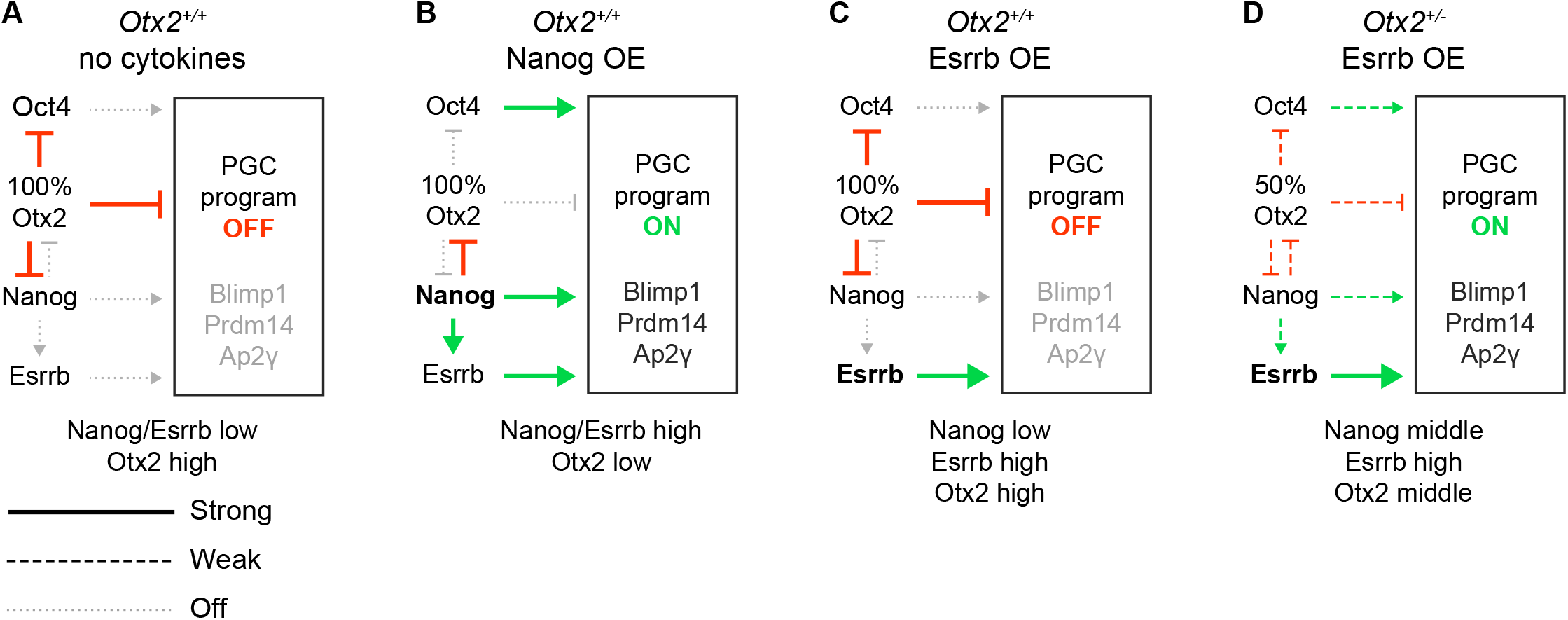
Proposed model of PGC induction. **A)** In wild-type EpiLCs, in the absence of BMP and associated cytokines, OTX2 levels remain above the threshold level required to block PGCLC differentiation during the period of germline competence. **B)** If NANOG expression is induced in EpiLCs at the start of differentiation, *Otx2* is repressed leading to a decreased OTX2 level that no longer represses expression of the OTX2 target genes *Nanog* and *Oct4* (Di Giovannantonio et al., 2021). This increases the expression of OCT4, which together with NANOG and its downstream target ESRRB provide positive regulatory inputs into the PGC GRN (centred on the TFs Blimp1, Prdm14 and Ap2γ), sufficient to drive PGCLC differentiation in the absence of cytokines. **C)** If ESRRB expression is induced, there is no repression of *Otx2* and the OTX2-mediated repression of *Nanog* and *Oct4* remain in place, blocking PGCLC differentiation. **D)** In *Otx2^+/-^* EpiLCs, repression of *Nanog* and *Oct4* by OTX2 is diminished in strength. This enables induction of ESRRB to provide sufficient input into the PGC GRN (alongside mild inputs from NANOG and OCT4) to direct PGCLC differentiation.

## Discussion

Despite discovery of the key transcription factors required for PGC identity, the rules governing PGC specification have yet to be fully elucidated. Both NANOG and ESRRB are important for maintaining PGC numbers *in vivo* (Chambers et al., 2007; Yamaguchi et al., 2009; M. Zhang et al., 2018) and NANOG is important for specifying germline entry *in vitro* (Murakami et al., 2016; J. Zhang et al., 2018; M. Zhang et al., 2018). During the first 24 hours of EpiLC – PGCLC differentiation *Otx2* is rapidly downregulated (J. Zhang et al., 2018). This downregulation is required to activate the PGC programme, since maintaining a high level of OTX2 during this time blocks PGCLC differentiation (J. Zhang et al., 2018). A key function of PGC-specifying cytokines during the early stages of EpiLC – PGCLC differentiation is to mediate this rapid *Otx2* downregulation (J. Zhang et al., 2018). In *Nanog*- null ESCs, *Otx2* is rapidly downregulated by NANOG induction and this is accompanied by H3K27me3 deposition at the *Otx2* promoter (Festuccia et al., 2012; Heurtier et al., 2019). In contrast, *Otx2* expression is induced during ESC – EpiLC transition and *Nanog* is repressed. Consequently, EpiLCs have high OTX2 levels and no *Nanog* expression (Acampora et al., 2017; Hayashi et al., 2011). In this study, we have examined the roles of the NANOG target genes *Esrrb* and *Otx2* during PGCLC specification.

The results reported here show that induction of NANOG in EpiLCs downregulates *Otx2* at the earliest time examined during differentiation and that this occurs in the absence of PGC-promoting cytokines. This further highlights similarities between the regulatory network of naïve pluripotent cells and early PGCs (Leitch and Smith, 2013). Importantly, enforced NANOG expression in EpiLCs induced a PGCLC population in the absence of the otherwise requisite cytokines (Murakami et al., 2016). The importance of the NANOG-mediated repression of *Otx2* for PGCLC differentiation under these conditions is demonstrated by the reduction in the size of the PGCLC population resulting from simultaneous enforced expression of both NANOG and OTX2. Therefore, OTX2 depletion is essential for PGCLC specification by NANOG in the absence of cytokines.

*Esrrb*, a direct target gene of NANOG, can substitute for NANOG in LIF-independent maintenance of mouse ESCs, reprogramming of pre-iPSCs / EpiSCs to naïve pluripotency (Festuccia et al., 2012). In addition, a knock-in of *Esrrb* to the *Nanog* locus can rescue the quantitative reduction in PGC number seen when *Nanog* is specifically deleted from the germ cell lineage (M. Zhang et al., 2018). Furthermore, the failure of *Nanog^-/-^* cells to maintain a PGCLC population in the presence of PGC-promoting cytokines could be rescued by enforced Esrrb expression (M. Zhang et al., 2018). It was somewhat surprising therefore, to see that ESRRB, unlike NANOG cannot efficiently induce PGC transcription factors *Blimp1* and *Prdm14* in the absence of cytokines. However, ESRRB overexpression in *Nanog^-/-^* ESCs does not fully recapitulate the effect of NANOG overexpression on ESC self-renewal (Festuccia et al., 2012). Although NANOG and ESRRB share common target genes (Festuccia et al., 2012), functional differences between NANOG and ESRRB exist since distinct subsets of genes are specifically regulated by either TF in ESCs (Sevilla et al., 2021). In addition, while the interactomes of NANOG and ESRRB contain common binding partners (Gagliardi et al., 2013; van den Berg et al., 2010), it is notable that ESRRB interacts with Mediator, suggesting that ESRRB may function prominently in transcriptional initiation. In contrast, the NANOG interactome lacks Mediator components (Gagliardi et al., 2013). While both interactomes link to NuRD and PcG, NANOG also links to Sin3a and NcoR complexes. These observations suggest that the balance between transcriptional activation and repression differs between NANOG and ESRRB, with NANOG showing more links to transcriptional repression than ESRRB. These distinctions are consistent with the prominent role proposed for transcriptional repression during PGCLC differentiation (Kurimoto et al., 2008). Consistent with this idea, we show here that the inability of enforced ESRRB expression to recapitulate the PGCLC differentiation induced by ectopic NANOG may be due to the inability of ESRRB to rapidly downregulate *Otx2* within the competence time window required for initiation of PGC differentiation. When we tested the effect of ESRRB induction in *Otx2^+/^* EpiLCs expressing ~50% of wild-type *Otx2* levels, ESRRB successfully activated expression of *Blimp1, Prdm14, Ap2γ* as well as the activity of the *Oct4* distal enhancer and surface expression of CD61/SSEA1. Therefore, the inability of ESRRB to induce the germline programme is overcome by a reduction in the level of OTX2 in the starting population.

In wild-type EpiLCs, PGC-promoting cytokines suppress *Otx2* expression and activate the PGC-specific gene regulatory network (GRN) incorporating *Blimp1, Prdm14* and *Ap2γ*, leading to effective PGCLC differentiation (J. Zhang et al., 2018). Our present findings on TF-driven PGCLC differentiation, conducted in the absence of PGC-promoting cytokines, can be incorporated into our current understanding of EpiLC-PGCLC differentiation as illustrated (Figure 4). In the absence of cytokines, OTX2 protein levels are sufficient to block activation of the PGCLC programme. Recent findings indicate that this repressive effect of OTX2 on the PCG programme results in large part from a direct repression of the *Nanog* and *Oct4* genes (Figure 4A) (Di Giovannantonio et al., 2021). Induction of transgenic Nanog in EpiLCs shifts the balance between the mutual antagonists NANOG and OTX2 in favour of NANOG (Figure 4B). Once OTX2 levels are sufficiently decreased this enables NANOG to act on regulatory elements controlling genes in the PGC-specific GRN, including *Blimp1, Prdm14* and *Ap2γ*as previously suggested (Murakami et al., 2016). In addition, NANOG can activate *Esrrb*, which may also positively affect the PGC-specific GRN. Moreover, a reduction in OTX2 eliminates suppression of *Oct4*, which can also positively feed into the PGC-specific GRN (Figure 4B). If instead, transgenic Esrrb is induced, a positive effect on the PGC-specific GRN can also be envisaged (Figure 4C). However, in this case, *Otx2* is not suppressed, OTX2 remains high, suppression of *Nanog* and *Oct4* is maintained and the positive input into the PGC-specific GRN is insufficient to activate PGCLC differentiation. In contrast, in *Otx2^+/-^* cells, the balance between the mutual antagonists OTX2 and NANOG shifts (Figure 4D). We suggest that in these conditions, induction of ESRRB provides a sufficient positive effect on the PGC-specific GRN to direct some PGC differentiation due to a weakened suppression of *Nanog* and *Oct4* by OTX2. Further work will be required to bring greater clarity to the early events involved in germline specification prior to activation of the PGC-specific GRN.

## Materials and Methods

### Cell culture and PGCLC differentiation

ESCs were routinely cultured in Glasgow Minimum Essential Medium (GMEM; Sigma, G5154) supplemented with 10% FCS (Gibco 10270-106), 100 U/ml LIF (homemade), Nonessential amino acids (Invitrogen, cat. 11140-036), 1 mM sodium pyruvate (Invitrogen, 11360-039), 2 mM L-glutamine (Invitrogen, 25030-024) and 50 nM 2-mercaptoethanol (Gibco 31350-010) at density of 3-10×10^5^ cells per cm^2^ in culture flasks coated with 0.1% gelatine (Smith, 1991). Presence of mycoplasma contamination was routinely tested.

EpiLC and PGCLC differentiation were performed as previously described (Hayashi and Saitou, 2013) with the difference that no cytokines were used for PGCLC specification from EpiLCs. ESCs were adapted to N2B27 medium supplemented with 0.4μM PD0325901, 3μM CHIR99021 and 100 U/ml LIF (homemade) for at least three passages. EpiLC state was induced by culturing 1×10^5^ ESCs on fibronectin coated plates in N2B27 medium supplemented with 20 ng/ml Activin A and 12 ng/ml bFGF for two days. EpiLCs were dissociated and aggregated in GK15 medium (15% knockout serum replacement, 1x nonessential amino acids, 2 mM L-glutamine, 1 mM sodium pyruvate, 1 mM 2-mercaptoehanol and 50 U/ml Penicillin/Streptomycin in GMEM) in the presence or absence of 1 μg/ml doxycycline. Aggregation was performed in 20 μl drops containing 2×10^3^ cells per drop hanging over PBS for two days at 37 °C. Aggregates were collected in 5 ml of GK15 medium with or without doxycycline and transferred to untreated culture plates. Aggregates were cultured for the next 4 days at 72 r.p.m at 37 °C and 7 % CO_2_.

### Generation of doxycycline inducible cell lines

To generate *Otx2^+/+^* Nanog or Esrrb inducible cell lines (TgiN, TgiE), firstly CAG-rtTAms2 was integrated into the *Rosa26* locus of E14Tg2a ESCs as described (Bressan et al., 2017). E14Tg2a-rtTAms2 cells were then transfected with piggyBac-TetON-Nanog-PGK-Hph-pA or piggyBac-TetON-Esrrb-PGK-Hph-pA together with pCMV-hyPBASE (Yusa et al., 2011) plasmids using lipofectamine 3000 (Thermofisher; L3000015).

To generate TgiNG and TgiNO cells, E14Tg2a-rtTAms2 cells were transfected with piggyBac-TetO-FRT5-TdTomato-2a-HygR-FRT3-tk (Festuccia et al., 2012) and selected for clones with robust TdTomato expression in presence of doxycycline and low levels of TdTomato in the absence of doxycycline. To integrate Nanog-T2A-GFP and Nanog-T2A-Otx2 cassettes, *Rosa26:* rtTAms2; E14Tg2a-TetON-TdTomato ESCs were transfected with pShuttle-TetON-FRT5-(Flag)_3_-Nanog-T2A-GFP-IRES-PURO-FRT3 or pShuttle-TetON-FRT5-(Flag)_3_-Nanog-T2A-Otx2-IRES-PURO-FRT3 together with pPGK-FLPobpa (Adgene #13793) using Lipofectamine 3000 (Thermofisher; L3000015). Cells resistant to puromycin and without TdTomato expression in presence of doxycycline were selected.

To generate *Otx2^+/-^* ΔPE::GFP ESCs with doxycycline inducible transgenes, *Otx2^lacZ/fl^* ΔPE::GFP c11 ESCs (J. Zhang et al., 2018) were transfected with piggyBac-TetON-Nanog-PGK-Hph-pA or piggyBac-TetON-Esrrb-PGK-Hph-pA together with pCMV-hyPBASE (Yusa et al., 2011) and pCAG-rTtAsm2-IRES-BSD using lipofectamine 3000 (Thermofisher; L3000015). Cells were selected with hygromycin and blasticidin for twelve days and single clones were picked. Two clones for Esrrb and one for Nanog were selected with capacity to induce the transgene and which express similar levels of Otx2 to the parental cell line.

### Flow cytometry

Cell aggregates were collected, washed with PBS and dissociated in 0.05 % Trypsin for ~10 minutes at 37 °C. Trypsin was neutralised with 10% FCS in GMEMβ and cells were collected by centrifugation (3 min at 300g). Cells were resuspended in 100 μl of GMEMβ medium supplemented with 10% FCS, 1:200 v/v Alexa Fluor 647 anti-mouse/human CD15 (SSEA-1) (Biolegend, 125608) and 1:500 v/v PE anti-mouse/rat CD61 (Biolegend, 104307). Cells were stained for 15 min at room temperature and washed twice with PBS before analysis at BD Fortessa 5 laser system.

### RT-qPCR

Single cell suspension was collected by centrifugation (3 min at 300 g) and total RNA was extracted by Illustra™ RNAspin RNA Isolation Kit (GE Healthcare GE25-0500-72) according to manufacturer instructions. First strand cDNA was synthetised using 200-1000 ng RNA, SuperScript III reverse transcriptase (Thermofisher 18080044) and random hexamers at the final concentration 2.5 mM. cDNA was diluted 1:10 in nuclease-free grade water. qPCR mix was prepared by mixing 5 μl cDNA, 4.5 μl of 2x Takyon SYBR Green master mix (Eurogentec, UF-NSMT-B0701) and 0.5 μl of 10 mM mixture of a primer pair (Supplementary table 1). Specificity and efficiency of used primer pairs were tested prior the experiments. Quantitative polymerase reactions were performed in two technical replicates for each sample using LightCycler 450 instrument (Roche).

### Data analysis

Flow cytometry data were analysed by FlowJo X 0.7 software. Single cells and DAPI negative cells were gated based on unstained and single stain controls. All other data analysis was done using R v4.0.3 (Team and others, 2013) and tidyverse packages (Wickham et al., 2019). Code and data to reproduce the analysis and figures are available at https://github.com/MatusV8/Otx2het.

---

**Supplementary Figure 1:**
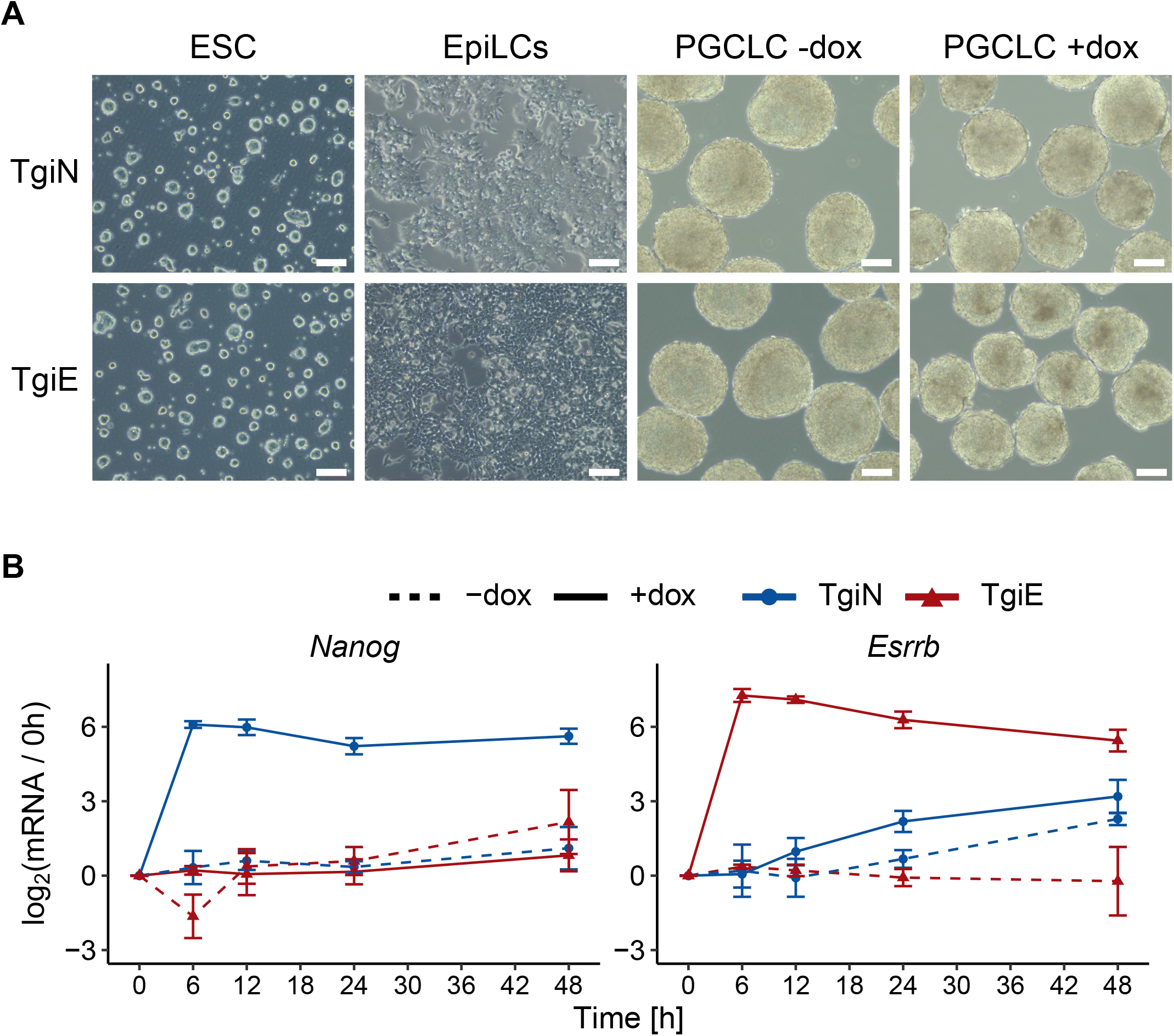
Preparation of doxycycline inducible Nanog and Esrrb cell lines. **A)** Photographs of TgiN and TgiE ESCs, EpiLCs and cell aggregates at day 6 of cytokine-free PGCLC differentiation in the presence (+dox) or absence (-dox) of doxycycline. White bars represent 100 μm. **B)** Relative changes of Esrrb and Nanog mRNAs after aggregation of E14Tg2a TetON-Nanog (TgiN) and E14Tg2a TetON-Esrrb (TgiE) EpiLCs cultured in the absence (-dox) or presence (+dox) of doxycycline at the indicated time point. Lines, points and triangles represent mean log2 fold-change (FC) differences between data points and the zero-time timepoint (mean ± SD, n = 3).

**Supplementary Figure 2:**
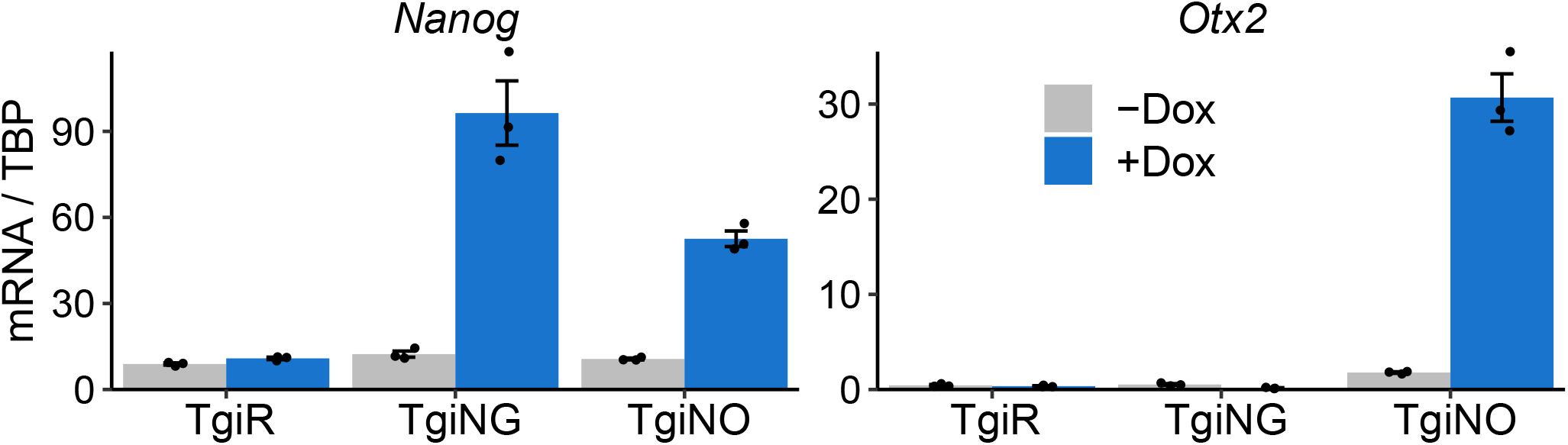
Activity of transgenes in TgiNG and TgiNO ESCs. RT-qPCR quantification of *Nanog* and *Otx2* mRNAs in TgiR, TgiNG and TgiNO ESCs (related to Figure 1A) before and after 48 hours of doxycycline treatment. Bar graph represents mean ± SEM (n=3) and the scatterplot represents individual data points.

**Supplementary Figure 3:**
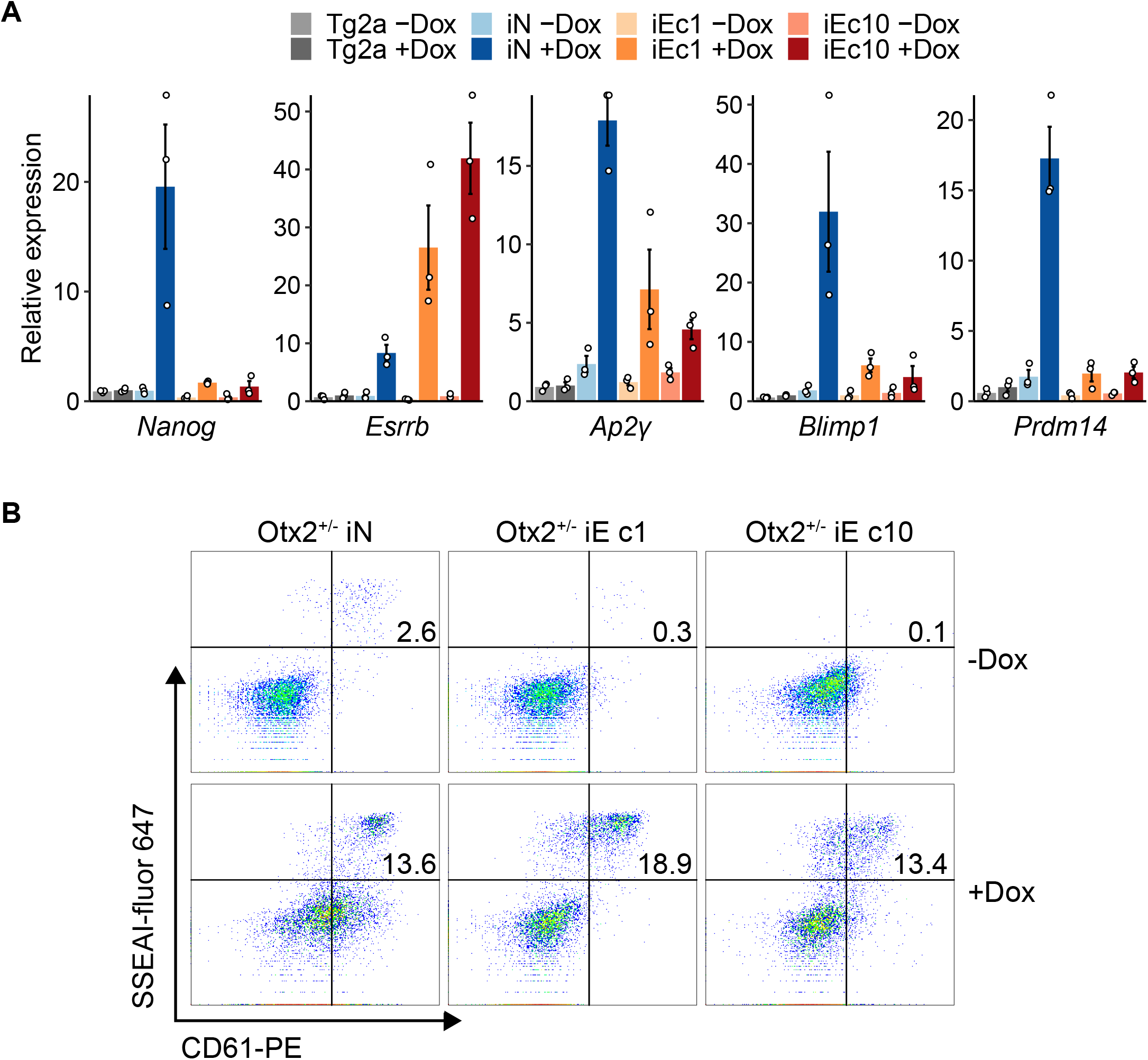
Cytokine-free differentiation of Nanog and Esrrb inducible *Otx2^+/-^* cell lines. **A)** RT-qPCR analysis of the indicated mRNAs at day 2 of the cytokine-free PGCLC differentiation in the presence (+Dox) or absence (-Dox) of doxycycline. Bars represent mean ± SEM (n = 3) of relative expression normalised to the mean expression in E14Tg2a -Dox sample for each mRNA. **B)** Representative flow cytometry analysis of SSEA1 and CD61 expression at day 6 of cytokine-free PGCLC differentiation of the indicated cell lines in the presence (+Dox) or absence (-Dox) of doxycycline. Percentages of SSEA1+CD61+ populations are indicated.

## Acknowledgements

We thank members of the Chambers lab for discussions and comments on the manuscript. This work was funded by grants to I.C. from the Biotechnological and Biological Sciences Research Council (BB/L002736/1) and the Medical Research Council (MR/T003162/1) of the UK and to M.Z. from the National Natural Science Foundation of China (Grant No. 32070800). M.V. was supported by a Principal’s Career Development Scholarship from University of Edinburgh.

## References

Acampora, D., Di Giovannantonio, L.G., Garofalo, A., Nigro, V., Omodei, D., Lombardi, A., Zhang, J., Chambers, I., Simeone, A., 2017. Functional Antagonism between OTX2 and NANOG Specifies a Spectrum of Heterogeneous Identities in Embryonic Stem Cells. Stem Cell Reports 9, 1642–1659. https://doi.org/10.1016/j.stemcr.2017.09.019

Acampora, D., Giovannantonio, L.G.D., Simeone, A., 2013. Otx2 is an intrinsic determinant of the embryonic stem cell state and is required for transition to a stable epiblast stem cell condition. Development 140, 43–55. https://doi.org/10.1242/dev.085290

Bressan, R.B., Dewari, P.S., Kalantzaki, M., Gangoso, E., Matjusaitis, M., Garcia-Diaz, C., Blin, C., Grant, V., Bulstrode, H., Gogolok, S., Skarnes, W.C., Pollard, S.M., 2017. Efficient CRISPR/Cas9-assisted gene targeting enables rapid and precise genetic manipulation of mammalian neural stem cells. Development 144, 635–648. https://doi.org/10.1242/dev.140855

Carter, A.C., Davis-Dusenbery, B.N., Koszka, K., Ichida, J.K., Eggan, K., 2014. Nanog-independent reprogramming to iPSCs with canonical factors. Stem Cell Reports 2, 119–126. https://doi.org/10.1016/j.stemcr.2013.12.010

Chambers, I., Silva, J., Colby, D., Nichols, J., Nijmeijer, B., Robertson, M., Vrana, J., Jones, K., Grotewold, L., Smith, A., 2007. Nanog safeguards pluripotency and mediates germline development. Nature 450, 1230–1234. https://doi.org/10.1038/nature06403

Di Giovannantonio, L.G., Acampora, D., Omodei, D., Nigro, V., Barba, P., Barbieri, E., Chambers, I., Simeone, A., 2021. Direct repression of Nanog and Oct4 by OTX2 modulates contribution of epiblast-derived cells to germline and somatic lineage. Development. https://doi.org/10.1242/dev.199166

Festuccia, N., Osorno, R., Halbritter, F., Karwacki-Neisius, V., Navarro, P., Colby, D., Wong, F., Yates, A., Tomlinson, S.R., Chambers, I., 2012. Esrrb Is a Direct Nanog Target Gene that Can Substitute for Nanog Function in Pluripotent Cells. Cell Stem Cell 11, 477–490. https://doi.org/10.1016/j.stem.2012.08.002

Gagliardi, A., Mullin, N.P., Ying Tan, Z., Colby, D., Kousa, A.I., Halbritter, F., Weiss, J.T., Felker, A., Bezstarosti, K., Favaro, R., Demmers, J., Nicolis, S.K., Tomlinson, S.R., Poot, R.A., Chambers, I., 2013. A direct physical interaction between Nanog and Sox2 regulates embryonic stem cell self-renewal. The EMBO Journal 32, 2231–2247. https://doi.org/10.1038/emboj.2013.161

Hayashi, K., Ohta, H., Kurimoto, K., Aramaki, S., Saitou, M., 2011. Reconstitution of the Mouse Germ Cell Specification Pathway in Culture by Pluripotent Stem Cells. Cell 146, 519–532. https://doi.org/10.1016/j.cell.2011.06.052

Hayashi, K., Saitou, M., 2013. Generation of eggs from mouse embryonic stem cells and induced pluripotent stem cells. Nature Protocols 8, 1513–1524. https://doi.org/10.1038/nprot.2013.090

Heurtier, V., Owens, N., Gonzalez, I., Mueller, F., Proux, C., Mornico, D., Clerc, P., Dubois, A., Navarro, P., 2019. The molecular logic of Nanog-induced self-renewal in mouse embryonic stem cells. Nat Commun 10, 1–15. https://doi.org/10.1038/s41467-019-09041-z

Kurimoto, K., Yabuta, Y., Ohinata, Y., Shigeta, M., Yamanaka, K., Saitou, M., 2008. Complex genome-wide transcription dynamics orchestrated by Blimp1 for the specification of the germ cell lineage in mice. Genes & Development 22, 1617–1635. https://doi.org/10.1101/gad.1649908

Lawson, K.A., Dunn, N.R., Roelen, B. a. J., Zeinstra, L.M., Davis, A.M., Wright, C.V.E., Korving, J., Hogan, B.L.M., 1999. Bmp4 is required for the generation of primordial germ cells in the mouse embryo. Genes & Development 13, 424–436. https://doi.org/10.1101/gad.13.4.424

Leitch, H.G., Smith, A., 2013. The mammalian germline as a pluripotency cycle. Development 140, 2495–2501. https://doi.org/10.1242/dev.091603

Magnusdottir, E., Dietmann, S., Murakami, K., Guenesdogan, U., Tang, F., Bao, S., Diamanti, E., Lao, K., Gottgens, B., Surani, M.A., 2013. A tripartite transcription factor network regulates primordial germ cell specification in mice. Nature Cell Biology 15, 905–U322. https://doi.org/10.1038/ncb2798

Mitsunaga, K., Araki, K., Mizusaki, H., Morohashi, K., Haruna, K., Nakagata, N., Giguère, V., Yamamura, K., Abe, K., 2004. Loss of PGC-specific expression of the orphan nuclear receptor ERR-β results in reduction of germ cell number in mouse embryos. Mechanisms of Development 121, 237–246. https://doi.org/10.1016/j.mod.2004.01.006

Murakami, K., Guenesdogan, U., Zylicz, J.J., Tang, W.W.C., Sengupta, R., Kobayashi, T., Kim, S., Butler, R., Dietmann, S., Surani, M.A., 2016. NANOG alone induces germ cells in primed epiblast in vitro by activation of enhancers. Nature 529, 403-+. https://doi.org/10.1038/nature16480

Ohinata, Y., Payer, B., O’Carroll, D., Ancelin, K., Ono, Y., Sano, M., Barton, S.C., Obukhanych, T., Nussenzweig, M., Tarakhovsky, A., Saitou, M., Surani, M.A., 2005. Blimp1 is a critical determinant of the germ cell lineage in mice. Nature 436, 207–213. https://doi.org/10.1038/nature03813

Sevilla, A., Papatsenko, D., Mazloom, A.R., Xu, H., Vasileva, A., Unwin, R.D., LeRoy, G., Chen, E.Y., Garrett-Bakelman, F.E., Lee, D.-F., Trinite, B., Webb, R.L., Wang, Z., Su, J., Gingold, J., Melnick, A., Garcia, B.A., Whetton, A.D., MacArthur, B.D., Ma’ayan, A., Lemischka, I.R., 2021. An Esrrb and Nanog Cell Fate Regulatory Module Controlled by Feed Forward Loop Interactions. Front. Cell Dev. Biol. 9. https://doi.org/10.3389/fcell.2021.630067

Smith, A.G., 1991. Culture and differentiation of embryonic stem cells. Journal of Tissue Culture Methods 13, 89–94. https://doi.org/10.1007/BF01666137

Team, R.C., others, 2013. R: A language and environment for statistical computing.

Urlinger, S., Baron, U., Thellmann, M., Hasan, M.T., Bujard, H., Hillen, W., 2000. Exploring the sequence space for tetracycline-dependent transcriptional activators: Novel mutations yield expanded range and sensitivity. Proc Natl Acad Sci U S A 97, 7963–7968.

van den Berg, D.L.C., Snoek, T., Mullin, N.P., Yates, A., Bezstarosti, K., Demmers, J., Chambers, I., Poot, R.A., 2010. An Oct4-centered protein interaction network in embryonic stem cells. Cell Stem Cell 6, 369–381. https://doi.org/10.1016/j.stem.2010.02.014

Vincent, S.D., Dunn, N.R., Sciammas, R., Shapiro-Shalef, M., Davis, M.M., Calame, K., Bikoff, E.K., Robertson, E.J., 2005. The zinc finger transcriptional repressor Blimp1/Prdm1 is dispensable for early axis formation but is required for specification of primordial germ cells in the mouse. Development 132, 1315–1325. https://doi.org/10.1242/dev.01711

Weber, S., Eckert, D., Nettersheim, D., Gillis, A.J.M., Schäfer, S., Kuckenberg, P., Ehlermann, J., Werling, U., Biermann, K., Looijenga, L.H.J., Schorle, H., 2010. Critical function of AP-2 gamma/TCFAP2C in mouse embryonic germ cell maintenance. Biol. Reprod. 82, 214–223. https://doi.org/10.1095/biolreprod.109.078717

Wickham, H., Averick, M., Bryan, J., Chang, W., McGowan, L.D., François, R., Grolemund, G., Hayes, A., Henry, L., Hester, J., Kuhn, M., Pedersen, T.L., Miller, E., Bache, S.M., Müller, K., Ooms, J., Robinson, D., Seidel, D.P., Spinu, V., Takahashi, K., Vaughan, D., Wilke, C., Woo, K., Yutani, H., 2019. Welcome to the Tidyverse. Journal of Open Source Software 4, 1686. https://doi.org/10.21105/joss.01686

Yamaguchi, S., Kurimoto, K., Yabuta, Y., Sasaki, H., Nakatsuji, N., Saitou, M., Tada, T., 2009. Conditional knockdown of Nanog induces apoptotic cell death in mouse migrating primordial germ cells. Development 136, 4011–4020. https://doi.org/10.1242/dev.041160

Yamaji, M., Seki, Y., Kurimoto, K., Yabuta, Y., Yuasa, M., Shigeta, M., Yamanaka, K., Ohinata, Y., Saitou, M., 2008. Critical function of Prdm14 for the establishment of the germ cell lineage in mice. Nature Genetics 40, 1016–1022. https://doi.org/10.1038/ng.186

Yusa, K., Zhou, L., Li, M.A., Bradley, A., Craig, N.L., 2011. A hyperactive piggyBac transposase for mammalian applications. Proc Natl Acad Sci U S A 108, 1531–1536. https://doi.org/10.1073/pnas.1008322108

Zhang, J., Zhang, M., Acampora, D., Vojtek, M., Yuan, D., Simeone, A., Chambers, I., 2018. OTX2 restricts entry to the mouse germline. Nature 562, 595–599. https://doi.org/10.1038/s41586-018-0581-5

Zhang, M., Chambers, I., 2019. Segregation of the mouse germline and soma. Cell Cycle 18, 3064–3071. https://doi.org/10.1080/15384101.2019.1672466

Zhang, M., Leitch, H.G., Tang, W.W.C., Festuccia, N., Hall-Ponsele, E., Nichols, J., Surani, M.A., Smith, A., Chambers, I., 2018. Esrrb Complementation Rescues Development of Nanog-Null Germ Cells. Cell Reports 22, 332–339. https://doi.org/10.1016/j.celrep.2017.12.060

